# The H3K9 methyltransferase SETDB1 maintains female identity in *Drosophila* germ cells by repressing expression of key spermatogenesis genes

**DOI:** 10.1101/259473

**Authors:** Anne E. Smolko, Laura Shapiro-Kulnane, Helen K. Salz

## Abstract

The preservation of germ cell sexual identity is essential for gametogenesis. Here we show that H3K9me3-mediated gene silencing is integral to female fate maintenance in *Drosophila* germ cells. Germ cell-specific loss of the H3K9me3 pathway members, the trimethyltransferase SETDB1, its binding partner WDE, and the H3K9 binding protein HP1a, cause the inappropriate expression of testis genes. SETDB1 is required for H3K9me3 accumulation on a select subset of the silenced testis genes. Interestingly, these SETDB1-dependent H3K9me3 domains are highly localized and do not spread into neighboring loci. Regional deposition is especially striking at the *phf7* locus, a key regulator of male germ cell sexual fate. *phf7* is primarily regulated by alternative promoter usage and transcription start site (TSS) selection. We find H3K9me3 accumulation is restricted to the silenced testis-specific TSS region in ovaries. Furthermore, its recruitment to *phf7* and repression of the testis-specific transcript is dependent on the female sex determination gene *Sxl*. These findings demonstrate that female identity is secured by a pathway in which *Sxl* is the upstream female-specific regulator, SETDB1 is the required chromatin writer and *phf7* is one of the critical SETDB1 target genes. This function of SETDB1 is unrelated to its canonical role in piRNA biogenesis and silencing of transposable elements. Collectively our findings support a novel model in which female fate is preserved by deposition of H3K9me3 repressive marks on key spermatogenesis genes and suggest that this strategy for securing cell fate may be widespread.

## Introduction

In metazoans, germ cell development begins early in embryogenesis when the primordial germ cells are specified as distinct from somatic cells. Specified primordial germ cells then migrate into the embryonic gonad, where they begin to exhibit sex-specific division rates and gene expression programs, ultimately leading to meiosis and differentiation into either eggs or sperm. Defects in sex-specific programming interferes with germ cell differentiation leading to infertility and germ cell tumors. Successful reproduction, therefore, depends on the capacity of germ cells to maintain their sexual identity in the form of sex-specific regulation of gene expression (Lesch and Page, 2012; Salz et al., 2017; Spiller et al., 2017).

In *Drosophila melanogaster*, germ cell sexual identity is specified in embryogenesis by the sex of the developing somatic gonad (Casper and van Doren, 2009; Horabin et al., 1995; Staab et al., 1996; Wawersik et al., 2005). However, extrinsic control is lost after embryogenesis and sexual identity is preserved by a cell-intrinsic mechanism (Casper and van Doren, 2009). The SEX-LETHAL (SXL) female-specific RNA binding protein is an integral component of this cell-intrinsic mechanism as loss of the protein specifically in germ cells leads to a global upregulation of spermatogenesis genes and a germ cell tumor phenotype (Chau et al., 2009; Schüpbach, 1985; Shapiro-Kulnane et al., 2015). Remarkably, sex-inappropriate transcription of a single gene, *PHD finger protein 7 (phf7)*, a key regulator of male identity (Yang et al., 2012), is largely responsible for the tumor phenotype (Shapiro-Kulnane et al., 2015). Depletion of *phf7* in these mutants suppresses the tumor phenotype and restores oogenesis. Moreover, forcing PHF7 protein expression in ovarian germ cells is sufficient to disrupt female fate and give rise to a germ cell tumor. Interestingly, sex-specific regulation of *phf7* is achieved by a mechanism that relies primarily on alternative promoter choice and transcription start site (TSS) selection. Sex-specific transcription produces mRNA isoforms with different 5’ untranslated regions that affect translation efficiency, such that PHF7 protein is only detectable in the male germline (Shapiro-Kulnane et al., 2015; Yang et al., 2012; Yang et al., 2017). Although the SXL protein is known to control expression post-transcriptionally in other contexts (Salz and Erickson, 2010), the observation that germ cells lacking SXL protein show defects in *phf7* transcription argues that *Sxl* is likely to indirectly control *phf7* promoter choice. Thus, how this sex-specific gene expression program is stably maintained remains to be determined.

Here, we report our discovery that female germ cell fate is maintained by an epigenetic regulatory pathway in which SETDB1 (aka EGGLESS, KMT1E and ESET) is the required chromatin writer and *phf7* is one of the critical SETDB1 target genes. SETDB1 trimethylates H3K9 (H3K9me3), a feature of heterochromatin (Brower-Toland et al., 2009; Elgin and Reuter, 2013). Using tissue-specific knockdown approaches we establish that germ cell specific loss of SETDB1, its protein partner WINDEI [WDE, aka ATF7IP, MCAF1 and hAM (Koch et al., 2009)], and the H3K9me3 reader, Heterochromatin binding protein 1a [HP1a, encoded by the *Su(var)205* locus (Eissenberg and Elgin, 2014)], leads to ectopic expression of euchromatic protein-encoding genes, many of which are normally expressed only in testis. We further find that H3K9me3 repressive marks accumulate in a SETDB1 dependent manner at 21 of these ectopically expressed genes, including *phf7*. Remarkably, SETDB1 dependent H3K9me3 deposition is highly localized and does not spread into neighboring loci. Regional deposition is especially striking at the *phf7* locus, where H3K9me3 accumulation is restricted to the region surrounding the silent testis-specific TSS. Lastly, we find that H3K9me3 accumulation at many of these genes, including *phf7*, is dependent on *Sxl*. Collectively our findings support a novel model in which female fate is preserved by deposition of H3K9me3 repressive marks on key spermatogenesis genes.

## Results

### H3K9me3 pathway members SETDB1, WDE and HP1a are required for germ cell differentiation

Previous studies reported that loss of the H3K9 trimethytransferase SETDB1 leads to mutant germaria filled with undifferentiated germ cells, characteristic of a germ cell tumor phenotype (Clough et al., 2007; Clough et al., 2014; Rangan et al., 2011; Wang et al., 2011; Yoon et al., 2008). Similarly, loss of WDE, SETDB1’s binding partner, also gives rise to a germ cell tumor (Koch et al., 2009). Because of the known connection between the germ cell tumor phenotype and ectopic testis gene transcription, we investigated the potential involvement of H3K9me3 marked chromatin in controlling expression of testis genes in female germ cells.

We first confirmed that the loss of SETDB1 and WDE specifically in germ cells was the cause of the germ cell tumor phenotype. To achieve <u>G</u>erm<u>L</u>ine specific <u>K</u>nock<u>D</u>own (*GLKD*), we expressed an inducible RNA interference (RNAi) transgene with *nos-Gal4*, a strong driver largely restricted to pre-meiotic germ cells (van Doren et al., 1998). As expected, *setdb1 GLKD* and *wde GLKD* abolished the intense H3K9me3 staining foci observed in wild-type germ cells (Fig. 1 A-C). We found the oogenesis defects elicited by *setdb1* and *wde GLKD* to be similar, as judged by the number of round spectrosome like structures present in the germarium (Fig. 1D). The spectrosome is a spherical a-Spectrin-containing structure that is normally found only in germline stem cells (GSCs) and its differentiating daughter cell, the cystoblast (~5 per germarium). As differentiation proceeds, the round spectrosome elongates and branches out to form a fusome. We find that the majority of *setdb1* and *wde GLKD* mutant germaria contain 6 or more spectrosome containing germ cells (Fig. 1E, F, H). This indicates that loss of SETDB1 and WDE in germ cells blocks differentiation, giving rise to a tumor phenotype.

**Fig. 1.**
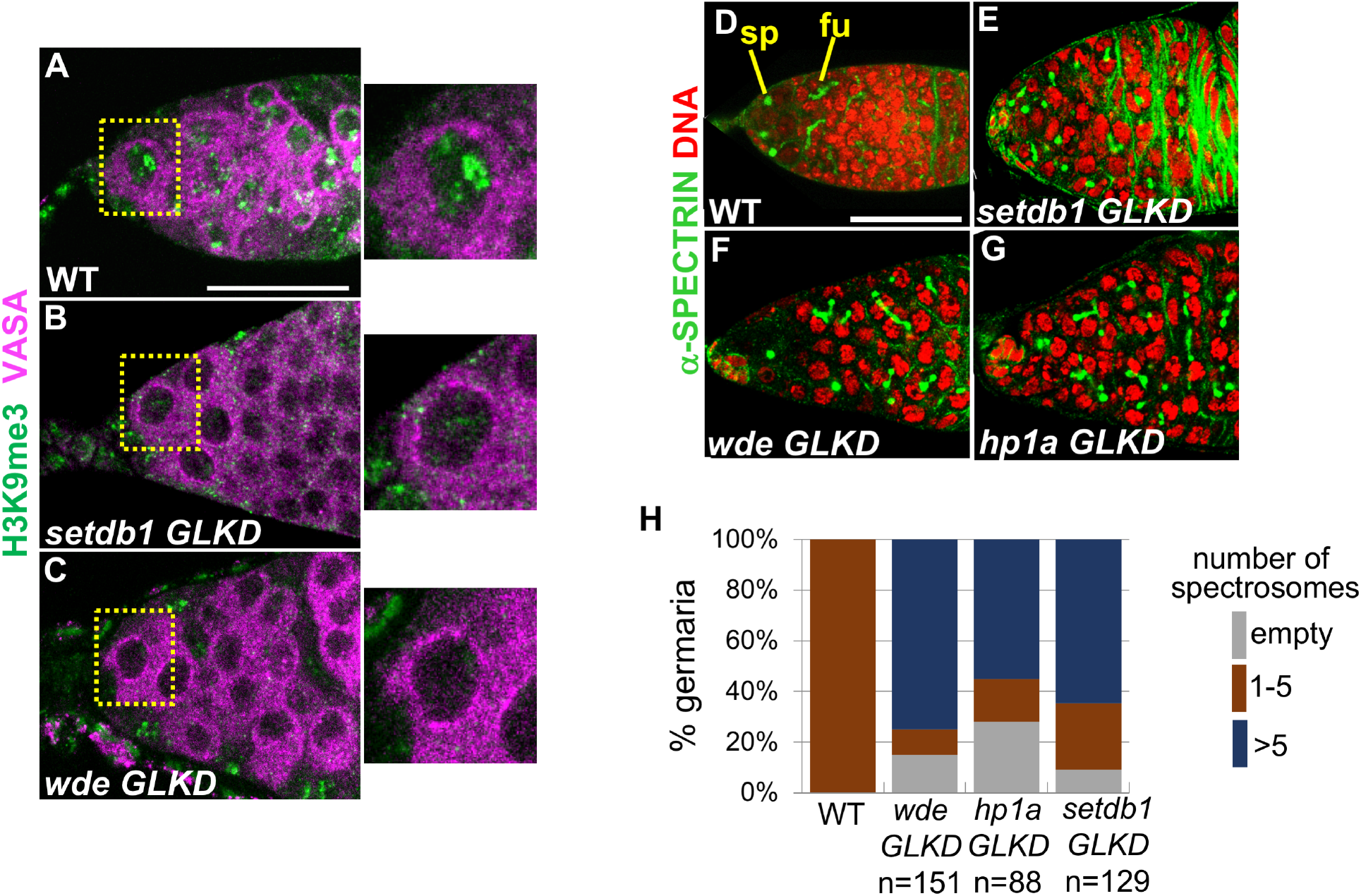
H3K9me3 is required for germ cell differentiation. **A-C** Reduced H3K9me3 staining in germ cells upon SETDB1 and WDE depletion. Representative confocal images of a wild-type (WT), *setdb1 GLKD, and wde GLKD* germarium stained for H3K9me3 (green). Germ cells were identified by α-VASA staining (magenta). Scale bar, 25 μm. Insets show a higher magnification of a single germ cell, outlined by a dashed line. **D-G** Undifferentiated germ cells accumulate in *setdb1 GLKD, wde GLKD* and *hp1a GLKD* mutant germaria. Representative confocal images of wild-type and mutant germaria stained for DNA (red) and α-spectrin (green) to visualize spectrosomes (sp), fusomes (fu) and somatic cells Scale bar, 25 μm. **H** Quantification of mutant germaria with 0, 1-5, and >5 round spectrosome-containing germ cells. The number of scored germaria (n) is indicated.

Recently, a large-scale RNAi screen identified a role for H3K9me3 binding protein HP1a in germ cell differentiation (Yan et al., 2014). In agreement with their findings, we observe that loss of the H3K9me3 binding protein HP1a in germ cells gives rise to germ cell tumors (Fig. 1G, H). HP1a binds to methylated H3K9 and is required for gene silencing in other contexts (Eissenberg and Elgin, 2014). This suggests that HP1a acts in a common pathway with SETDB1 and WDE in female germ cells.

### Loss of H3K9me3 pathway members in female germ cells leads to ectopic testis gene expression

To investigate the possibility that the loss of H3K9me3 pathway members in female germ cells leads to ectopic testis gene expression, we first used RT-qPCR to assay the *phf7* RNA isoforms present in mutant ovaries (Fig. 2A). Using a primer pair capable of detecting the male-specific *phf7-RC* transcript, we find that the testis specific *phf7-RC* transcript is ectopically expressed in *setdb1, wde*, and *hp1a GLKD* mutant ovaries. We therefore conclude that the H3K9me3 pathway members are essential for suppression of the male-specific *phf7* transcription in female germ cells.

**Fig. 2.**
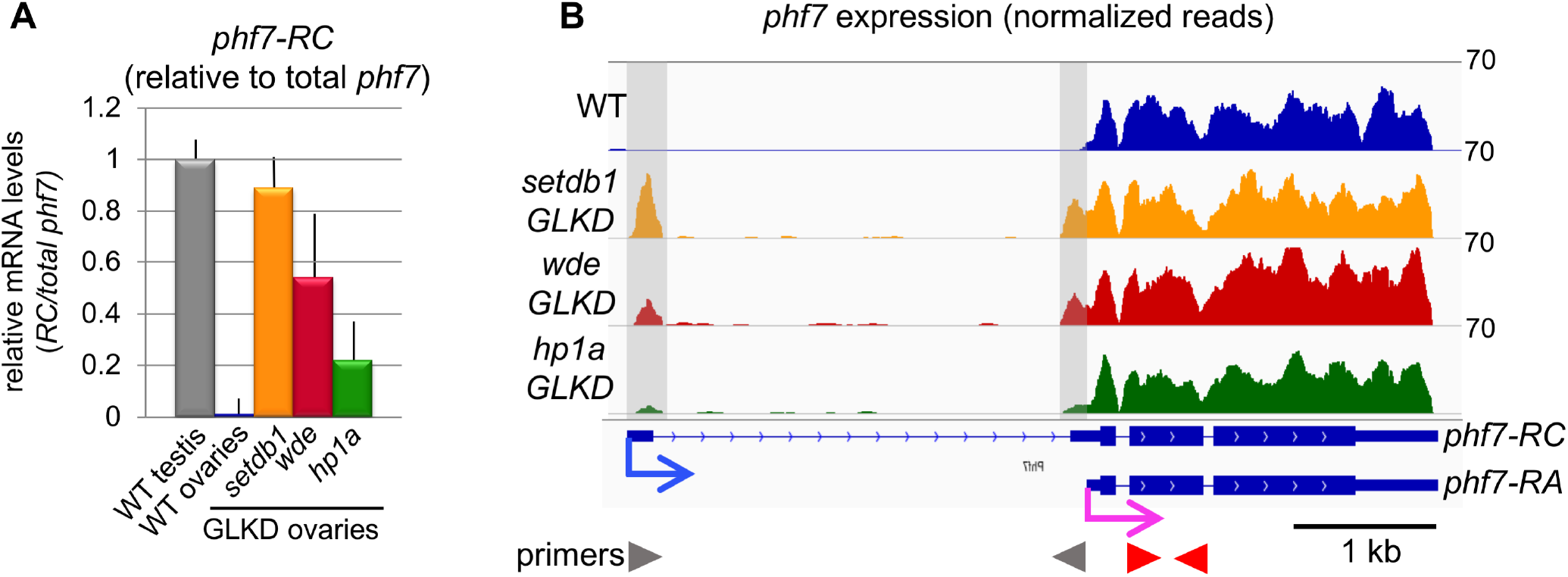
SETDB1, WDE, and HP1a depletion leads to female-to-male reprogramming at the *phf7* locus. **A** Depletion of H3K9me3 pathway members leads to ectopic expression of the testis-specific *phf7-RC* isoform. RT-qPCR analysis of the testis *phf7-RC* transcript in wild-type testis, wild-type and mutant ovaries. Expression, normalized to the total level of *phf7*, is shown as fold change relative to testis. Primers are shown in panel B. Error bars indicate SD of three biological replicates. **B** RNA-seq data confirms ectopic expression of the testis-specific *phf7-RC* isoform. Genome browser view of the *phf7* locus. Tracks show RNA-seq reads aligned to the *Drosophila* genome (UCSC dm6). All tracks are viewed at the same scale. The screen shot is reversed so that the 5’ end of the gene is on the left. The reads that are unique to the mutant *GLKD* ovaries are highlighted by gray shading. Beneath is the RefSeq gene annotation are the two *phf7* transcripts, *phf7-RC* and *phf7-RA*. Transcription start sites are indicated by blue (*phf7-RC*) and pink (*phf7-RA*) arrows. Primers for RT-qPCR are indicated by arrowheads: gray for *phf7-RC*, red for total *phf7*.

To gain a genome-wide view of the expression changes associated with the loss of H3K9me3 pathway members in germ cells, we used high throughput sequencing (RNA-seq) to compare the transcriptomes of *GLKD* mutant ovaries with wild-type ovaries from newborn (0-24 hour) females. New born ovaries lack late stage egg chambers, establishing a natural method for eliminating cell types not present in tumors. This minimizes the identification of gene expression changes unrelated to the mutant phenotype. From this analysis we identified 1191 genes in *setdb1* GLKD mutant ovaries, 904 in *wde* GLKD ovaries, and 1352 in *hp1a* GLKD ovaries that are upregulated at least 2-fold relative to wild-type (FDR<0.05; Tables S1-S3). Additionally, 657 genes in *setdb1* GLKD mutant ovaries, 756 in *wde* GLKD ovaries, and 877 in *hp1a* GLKD ovaries are downregulated (Tables S4-S6). Comparison of the differential gene expression profiles of *setdb1 GLKD* mutant ovaries with *wde* and *hpa1 GLKD* mutant ovaries revealed extensive similarities, as expected for genes functioning in the same pathway (Fig. 3A).

**Fig. 3.**
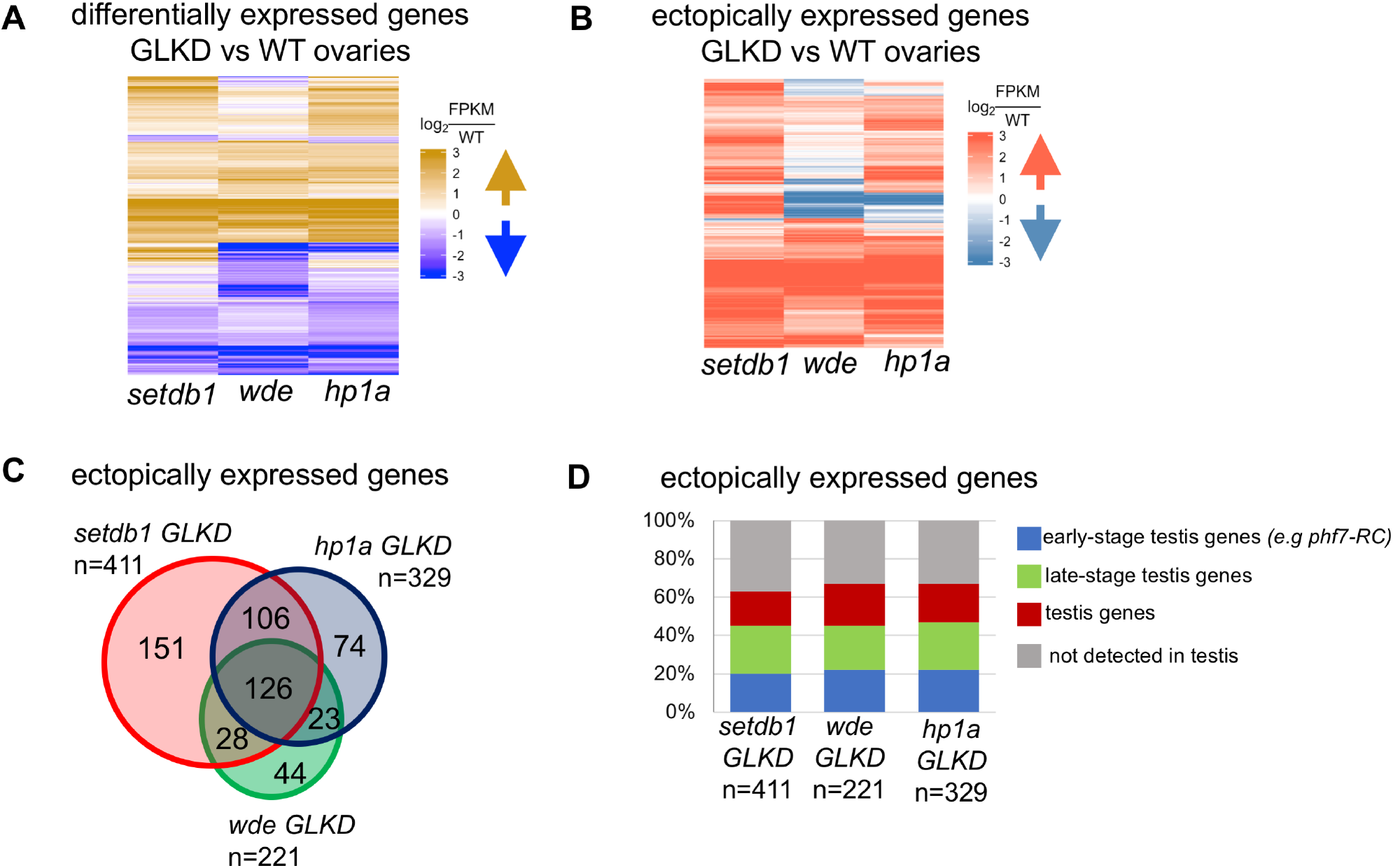
Depletion of H3K9me3 pathway members leads to ectopic expression of a similar set of genes normally expressed in the testis. **A** Transcriptome analysis by RNAs-seq shows that depletion of pathway members leads to a similar set of differentially expressed genes. Heat map comparing changes in gene expression in *setdb1, wde*, and *hp1a GLKD* ovaries compared to wild-type ovaries. Each row depicts a gene that changed expressed at least 2-fold (FDR<0.05) in at least one mutant when compared to wild-type. **B** Depletion of pathway members leads to a similar set of ectopically expressed genes. Heat map comparing ectopically expressed genes in *setdb1, wde*, and *hp1a GLKD* ovaries (FPKM < 1 in wild-type ovaries). Each row depicts a gene that is ectopically expressed in at least one mutant. **C** Venn diagram showing overlap of ectopically expressed genes in *setdb1, wde*, and *hp1a GLKD* ovaries. The amount of overlap is significantly higher than expected stochastically (P<10^-6^). The significance of each two-way overlap was assessed using Fisher’s exact test performed in R, in each case yielding P<10^-6^. The significance of the three-way overlap was assessed by a Monte Carlo simulation, yielding P<10^-6^. For each genotype, the number (n) of ectopically expressed genes is indicated. **D** A majority of ectopically expressed genes are normally expressed in testis. Bar chart showing the percentage of ectopically expressed genes in *setdb1, wde*, and *hp1a GLKD* ovaries that are normally expressed in wild-type testis. Genes are assigned into groups based on expression in WT and *bgcn* mutant testis (see text for details): early stage spermatogonia genes (>2-fold increase in *bgcn* compared to WT, in blue), late-stage testis genes (>2-fold decrease in *bgcn* compared to WT, in green) and unassigned testis genes (FPKM>1 in either sample, in red). For each genotype, the number (n) of ectopically expressed genes is indicated.

In agreement with our RT-qPCR analysis, we find that the testis-specific *phf7* transcript, *phf7-RC*, is ectopically expressed in *setdb1, wde*, and *hp1a GLKD* mutant ovaries (Fig. 2B). Given the pivotal role of *phf7* in controlling germ cell sexual identity, this finding suggests that the upregulated genes might include anomalously expressed testis genes. Indeed, all mutants express a set of similarly upregulated genes that are not detected in wild-type ovaries (FPKM<1 in wild-type ovaries; Fig. 3B, C; Tables S7-S9).

Previous work has shown that male-specific *phf7* expression is restricted to undifferentiated germ cells (Yang et al., 2012; Yang et al., 2017). To determine whether the testis genes expressed in mutant ovaries are similarly stage-specific, we compared our data with the published RNA-seq analysis of wild-type testis and *bgcn* mutant testis (Shan et al., 2017). The spermatogenesis gene expression program includes a major change in transcriptional output when the undifferentiated mitotically active spermatogonia stop dividing and transition into differentiating spermatocytes (White-Cooper, 2010). In spermatogenesis, *bgcn* is required for the undifferentiated spermatogonia to stop mitosis and transition into the spermatocyte stage. In *bgcn* mutants this transition is blocked, and the testis are enriched for dividing spermatogonial cells. The comparison of the wild-type and mutant expression profiles can therefore be used to identify genes preferentially expressed in early-stage spermatogonia (>2-fold increase in *bgcn* compared to wild-type, in blue) and in late-stage spermatocytes (>2-fold decrease in *bgcn* compared to wild type, in green). We also identified genes that are normally expressed in testis but are not differentially expressed (FPKM > 1 in both samples, in red) and genes that are not detectable in either sample (FPKM <1, in gray). This analysis revealed that 63%-67% of the ectopically expressed genes are expressed in testis (Fig. 3D; Tables S7-S9). Because we identify examples of both early and late stage testis genes amongst the ectopically expressed genes, we surmise that the mutant germ cells are attempting to engage the spermatogenesis differentiation program. Together, these studies indicate that SETDB1-controlled H3K9 methylation secures female germ cell identity by silencing testis genes.

### The presence of SETDB1-dependent H3K9me3 islands at select genes correlates with sex-specific transcriptional regulation

Our studies raise the possibility that SETDB1 silences testis transcription by mediating the deposition of H3K9me3 on its target loci. To test this idea directly, we performed H3K9me3 chromatin immunoprecipitation followed by sequencing (ChIP-seq) on wild-type and *setdb1 GLKD* ovaries. By limiting the differential peak analysis to euchromatin genes, we identified 746 H3K9me3 enrichment peaks in wild-type that were significantly altered in *setdb1 GLKD* ovaries (Fig. 4A). Whereas a majority of the gene associated peaks show the expected decrease in H3K9me3 enrichment (84%, 630/746), we also observed regions with an increase in H3K9me3 enrichment (15%, 116/746). How loss of SETDB1 might lead to an increase in H3K9me3 is not known, but the effect is most likely indirect.

**Fig. 4.**
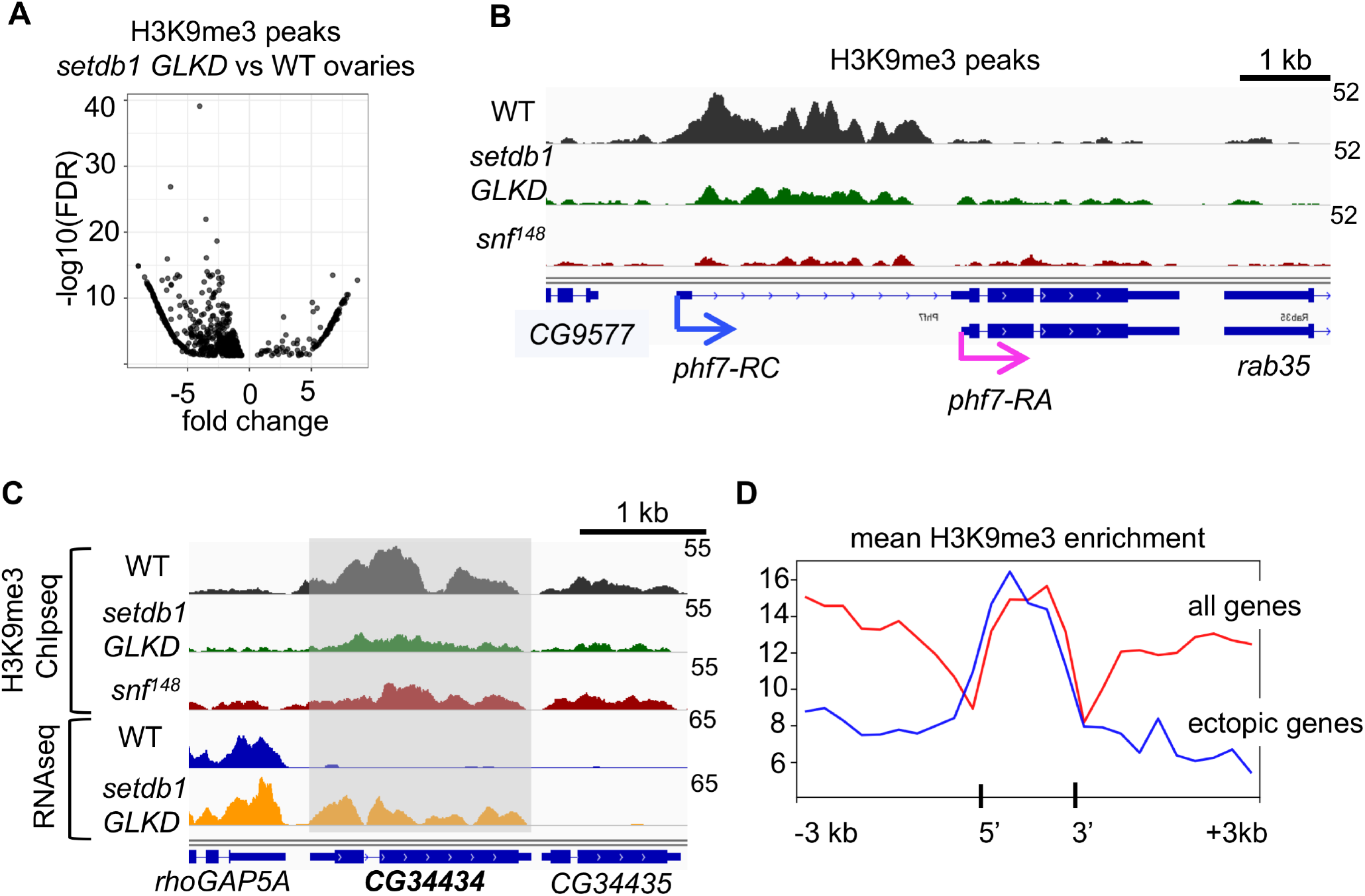
Loss of SETDB1 in female germ cells leads to H3K9me3 depletion on a select set of genes. **A** Differential analysis of paired H3K9me3 ChIP-seq data sets identifies SETDB1-dependent H3K9me3 peaks. Scatter plot showing the significantly altered (FDR < 5%) H3K9me3 peaks in *setdb1 GLKD* ovaries relative to wild-type (WT) ovaries. Negative values indicate a reduction in H3K9me3 in mutant ovaries. **B** The H3K9me3 peak over the testis-specific *phf7-RC* TSS is decreased in *setdb1 GLKD* and *snf^148^* mutant ovaries. Genome browser view of *phf7* and neighboring genes *CG9577* and *rab35*. **C** In *setdb1* GLKD, ectopic expression of *CG34435* is correlated with a decreased H3K9me3 ChIP-seq peak. This peak is also decreased in *snf^148^* mutant ovaries. Genome browser view of *CG34435* and its neighbors, *rhoGAP5A* and *CG34435*. RNA reads in wild-type (WT) and *setdb1 GLKD* are in blue and orange respectively. The ChIP-seq reads are in black, green and red. CG34435 is shaded. **D** Average enrichment profile indicates that the H3K9me3 peaks at the 21 SETDB1-regulated genes are localized over the gene body. The average H3K9me3 enrichment profile on the average gene body (transcription start site to transcription end site) scaled to 1500 base pairs, ± 3 kb. In red, the average enrichment profile of euchromatic genes which display an H3K9me3 peak in wild-type ovaries. In blue, the average H3K9me3 profile of the 21 genes which are both ectopically expressed and display a loss of H3K9me3 enrichment in *setdb1 GLKD* ovaries.

At the *phf7* locus, H3K9me3 is concentrated over the silenced testis-specific TSS in wild-type ovaries (Fig. 4B). Loss of this peak in *setdb1 GLKD* ovaries correlates with aberrant testis-specific transcription (Fig. 2B). These data suggest a functional link between the presence of repressive H3K9me3 chromatin and the silencing of *phf7* testis-specific isoform transcription.

We identified an additional 24 normally silenced euchromatic genes at which the loss of the H3K9me3 peak in *setdb1 GLKD* ovaries correlates with ectopic expression. Of these genes, 4 contain transposable element (TE) sequences. Prior studies have shown that H3K9me3 is enriched around euchromatic TE insertion sites, suggesting the possibility that transcriptional repression might result from spreading of H3K9me3 from a silenced TE (Lee, 2015; Lee and Karpen, 2017; Sienski et al., 2012). The absence of TE sequences at *phf7* and the other 20 genes suggests that H3K9me3 deposition is controlled by a novel mechanism. Thus, we will focus on these 21 genes (Table 1). Like *phf7*, the majority of these 20 SETDB1-regulated genes are normally expressed in testis. This gene set includes genes normally enriched in both early-stage germ cells and late-stage spermatocytes. Furthermore, an examination of their expression pattern in adult tissues, as reported in published data sets (Leader et al., 2018), indicates that 8 of these genes express testis-specific mRNAs. 7 are normally expressed in testis, but also in other tissues. The remaining 5 are not detectable in adult testis. Further studies are needed to determine whether sex-specific repression of these genes is as important for female germ cell development as *phf7*.

**Table 1.** SETDB1/H3K9me3 regulated genes

| **Gene** | **Protein domain, group membership and/or predicted function** | **RNA expression pattern in adult tissues** |
| --- | --- | --- |
| **TESTIS-SPECIFIC** |  |  |
| phf7 | H3K4me2 binding | early-stage |
| skpE | SKP1 gene family | early-stage |
| CG12477 | Ring finger domain/E3 ligase | early-stage |
| CG42299 | Zinc finger MIZ-type/E3 ligase | both |
| CG12061 | Sodium/calcium exchanger | both |
| CG13423 | Peptidase | late-stage |
| CG15172 | Unknown | late-stage |
| CG34434 | Unknown | late-stage |
| CR43299 | ncRNA | late-stage |
| **TESTIS-ENRICHED RELATIVE TO OVARIES** |  |  |
| Lim1 | Homeobox transcription factor | early-stage |
| CG32506 | Rab GTPase activating proteins | early-stage |
| CG42613 | CUB domain | early-stage |
| CG10440 | BTB/POZ domain | late-stage |
| CG10483 | SPX and EXS domains | late-stage |
| CG17636 | Hydrolase | late-stage |
| CG31202 | Alpha-mannosidase class I | late-stage |
| **NOT DETECTED IN TESTIS** |  |  |
| CG12607 | Unknown | no expression in adults |
| CG32679 | Secretory protein | no expression in adults |
| CG15818 | Carbohydrate binding | gut |
| MsR1 | Transmembrane G protein coupled receptor | brain/CNS |
| Rab3-GEF | GDP-GTP exchange factor | brain/CNS |

Co-regulated genes are often clustered together in the genome. In Drosophila, about a third of the testis-specific genes are found in groups of three or more genes (Parisi et al., 2004; Shevelyov et al., 2009). However, the 21 SETDB1-regulated genes we have identified do not fall within the previously identified testis-specific gene clusters, nor are they clustered together in the genome (Fig. 5, Table S10). Even on the X chromosome, where 11 of the 21 genes are located, the closest two genes are 100 kb away from each other. Thus, the SETDB1 regulated genes are not located within coexpression domains.

**Fig. 5.**
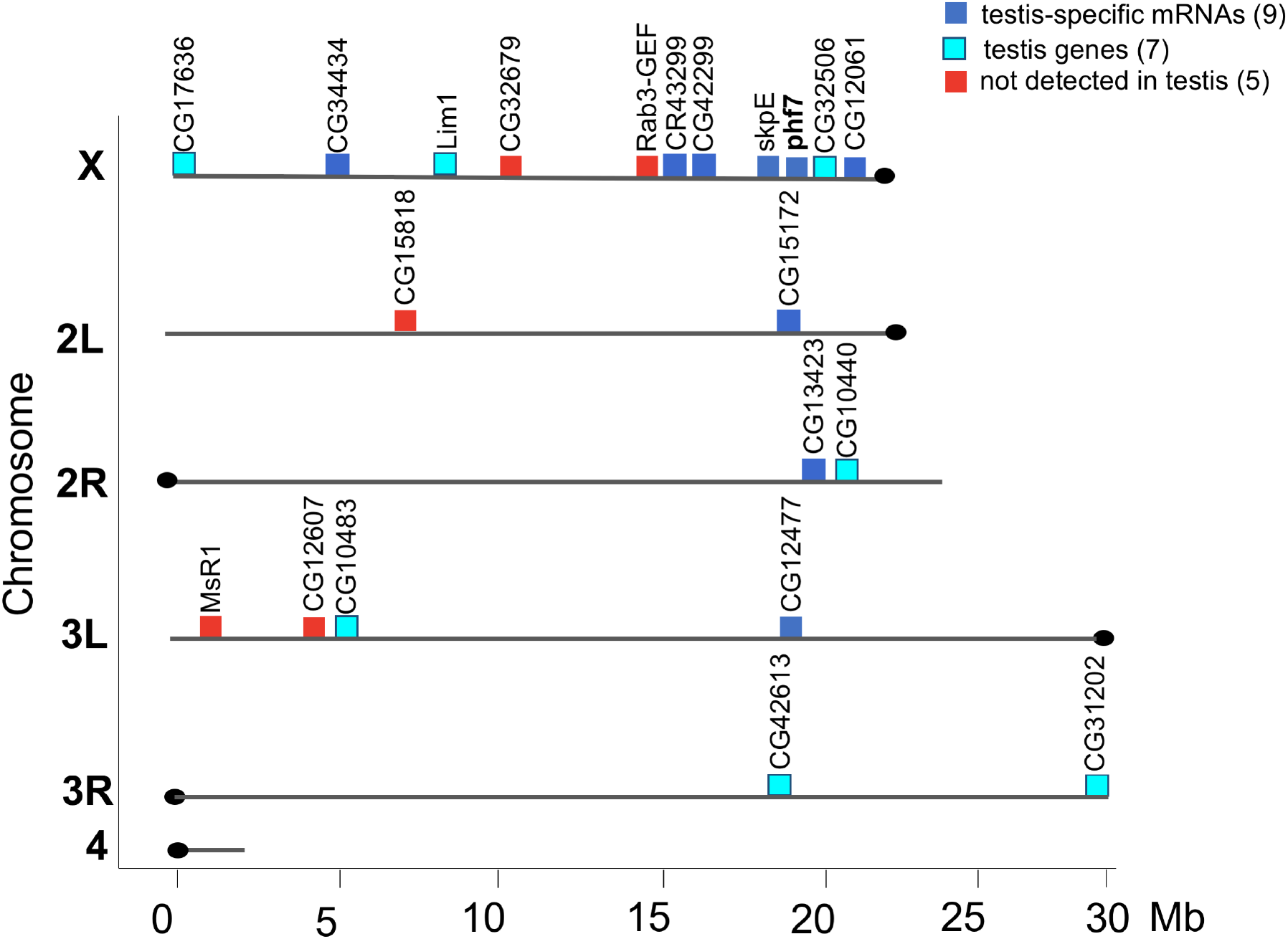
Distribution of the 21 SETDB1/H3K9me3 regulated genes in the genome. The 21 genes are not clustered together in the genome. Gene positions are shown on the six major chromosome arms. Chromosome length is indicated in megabase pairs (Mb). See Table S10 for the exact position of each gene. The normal expression pattern of these genes is indicated as follows: genes with testis-specific isoform (dark blue), genes expressed in testis and other tissues (turquoise) and genes not normally expressed in adult testis (red). See Table 1 for details.

In fact, the H3K9me3 peaks at the 21 SETDB1-regulated genes are highly localized and do not spread into the neighboring genes (*e.g*. Fig. 4B, C and Fig. S1). Averaging the H3K9me3 distribution over these 21 genes, scaled to 1.5 kb and aligned at their 5’ and 3’ ends, demonstrates a prominent enrichment over the gene body (Fig. 4D, in blue). In contrast, the average enrichment profile over all euchromatic genes with H3K9me3 peaks showed a broader pattern extending both upstream and downstream of the gene body (Fig. 4D, in red). Together these results clearly show that silencing involves formation of gene-specific blocks of H3K9me3 islands at a select set of testis genes.

### Loss of SXL protein in germ cells interferes with H3K9me3 accumulation

Previous studies have established that sex-specific *phf7* transcription is controlled by the female sex determination gene *Sxl* (Shapiro-Kulnane et al., 2015). Together with studies demonstrating that SXL protein is expressed in *setdb1* mutant germ cells (Clough et al., 2014), these data suggest that SXL acts upstream, or in parallel to SETDB1 to control *phf7* transcription. To assess the potential of a *Sxl*-mediated mechanism, we asked whether the loss of SXL in germ cells affects H3K9me3 accumulation. As with our earlier studies, we take advantage of the viable *sans-fille^148^ (snf^148^)* allele to selectively eliminate SXL protein in germ cells without disrupting function in the surrounding somatic cells (Chau et al., 2009; Chau et al., 2012; Shapiro-Kulnane et al., 2015). SNF, a general splicing factor, is essential for *Sxl* autoregulatory splicing (Salz and Erickson, 2010). The viable *snf^148^* allele specifically disrupts the *Sxl* autoregulatory splicing loop in female germ cells, leading to a failure in SXL protein accumulation and a germ cell tumor phenotype (Chau et al., 2009; Nagengast et al., 2003). We find that that the intense H3K9me3 staining foci observed in wild-type germ cells is reduced in *snf^148^* mutants (Fig. 6A). However, we find that SETDB1 protein expression, measured by an HA tag knocked into the endogenous locus (Seum et al., 2007), is not disrupted in *snf^148^* mutant germ cells (Fig. 6B). In both wild-type and *snf^148^* mutant germ cells, we find SETDB1 showed diffuse cytoplasmic staining and punctate nuclear staining (arrow head). Our finding that H3K9me3 staining is disrupted, even though SETDB1 protein accumulation appears normal in *snf^148^* mutant ovaries, leads us to conclude that SXL and SETDB1 collaboratively promote H3K9me3-mediated silencing (Fig. 6C).

**Fig. 6.**
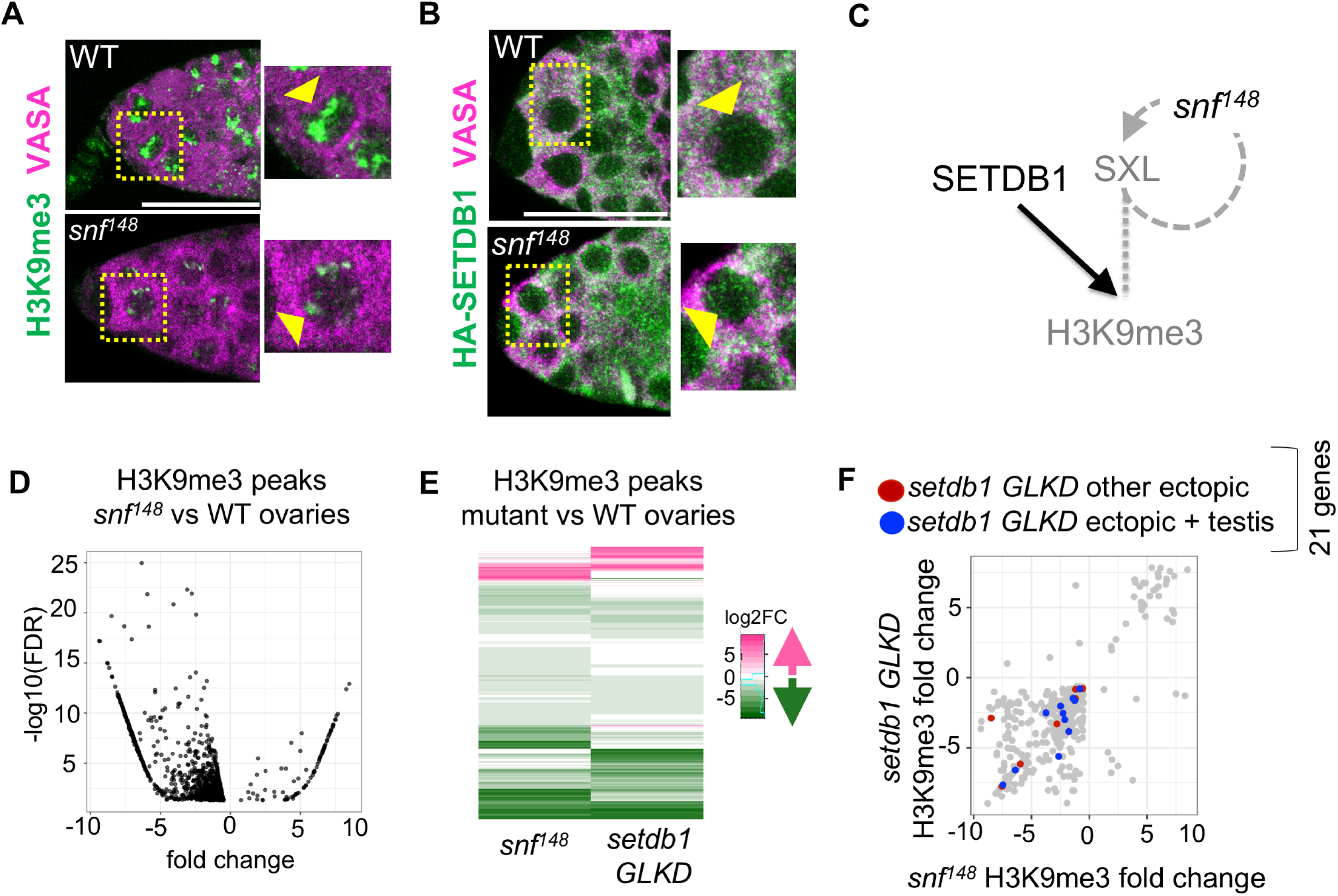
H3K9me3 distribution is affected in *snf^148^* mutant ovaries. **A** Reduced H3K9me3 staining in *snf^148^* mutant germ cells. Representative confocal images of a wild-type (WT) and *snf^148^* germaria stained for H3K9me3 (green). Germ cells were identified by α-VASA staining (magenta). Scale bar, 25 μm. Insets show a higher magnification of a single germ cell, outlined by a dashed line. **B** SETDB1 protein localization is not altered in *snf^148^* mutant germ cells. Representative confocal images of a germaria from wild-type and *snf^148^* females carrying a copy an endogenously HA-tagged allele of *setdb1* stained for HA (green). Germ cells were identified by α-VASA staining (magenta). Scale bar, 12.5 μm. Insets show a higher magnification of a single germ cell, outlined by a dashed line. **C** Diagram of genetic pathway controlling H3K9me3 accumulation in female germ cells. In WT, SXL collaborates with SETDB1 to regulated H3K9me3 accumulation. *snf^148^*, by virtue of the fact that it interferes with *Sxl* splicing, leads to germ cells without SXL protein. This in turn leads to a defect in H3K9me3 accumulation without interfering with SETDB1 protein accumulation. **D** Differential analysis of paired H3K9me3 ChIP-seq data sets identifies peak changes in *snf^148^* mutant ovaries. Scatter plot of significantly altered (FDR < 5%) H3K9me3 peaks in *snf^148^* ovaries relative to wild-type (WT) ovaries. Negative values indicate a decreased H3K9me3 peak in mutant ovaries. **E** *setdb1* GLKD and *snf^148^* mutant ovaries exhibit similar changes in H3K9me3 accumulation. Heat map comparing the differential H3K9me3 peak analysis carried out in *snf^148^* and *setdb1 GLKD* ovaries Each row depicts a gene with a significantly altered peak in either *snf^148^* or *setdb1 GLKD* ovaries compared to wild-type. **F** *setdb1 GLKD* and *snf^148^* influence H3K9me3 accumulation on a similar set of genes, including the genes ectopically expressed in *setdb1* GLKD ovaries. Plot comparing the significantly altered H3K9me3 peaks observed in *snf^148^* ovaries to *setdb1 GLKD* ovaries. Genes ectopically expressed in *setdb1 GLKD* are labeled in red while those that are ectopically expressed and normally expressed in testis are labeled in blue.

To directly test whether *Sxl* plays a role in controlling H3K9me3 deposition, we profiled the distribution of H3K9me3 by ChIP-seq in *snf^148^* mutant ovaries and compared it to the distribution in wild-type ovaries. By limiting the differential peak analysis to within 1 kb of euchromatic genes, we identified 1,039 enrichment peaks in wild-type that were significantly altered in *snf^148^* mutant ovaries, 91% of which show the expected decrease in H3K9me3 enrichment (Fig. 6D). The strong overlap between the regions displaying decreased H3K9me3 enrichment in *snf^148^* and *setdb1 GLKD* mutant ovaries suggests that SXL and SETDB1 influence H3K9me3 accumulation on the same set of genes (Fig. 6E, F), including *phf7* (Fig. 4B) and CG34434 (Fig. 4C). Based on these studies we conclude that SXL functions with SETDB1 in the assembly of H3K9me3 silencing islands at a select subset of sex-specific regulated genes.

## Discussion

This study reveals a novel role for H3K9me3 chromatin, operationally defined as facultative heterochromatin, in securing female identity by silencing lineage-inappropriate transcription. We show that H3K9me3 pathway members, the H3K9 methyltransferase SETDB1, its binding partner WDE, and the H3K9 binding protein HP1a, are required for silencing testis gene transcription in female germ cells. Our studies further suggest a mechanism in which SETDB1, in conjunction with the female fate determinant SXL, controls transcription through deposition of highly localized H3K9me3 islands on a select subset of testis genes. The male germ cell sexual identity gene *phf7* is one of the key downstream SETDB1 target genes. H3K9me3 deposition on the region surrounding the testis-specific TSS guaranties that no PHF7 protein is produced in female germ cells. In this model, failure to establish silencing leads to ectopic PHF7 protein expression, which in turn drives aberrant expression of testis genes and a tumor phenotype (Fig. 7).

**Fig. 7.**
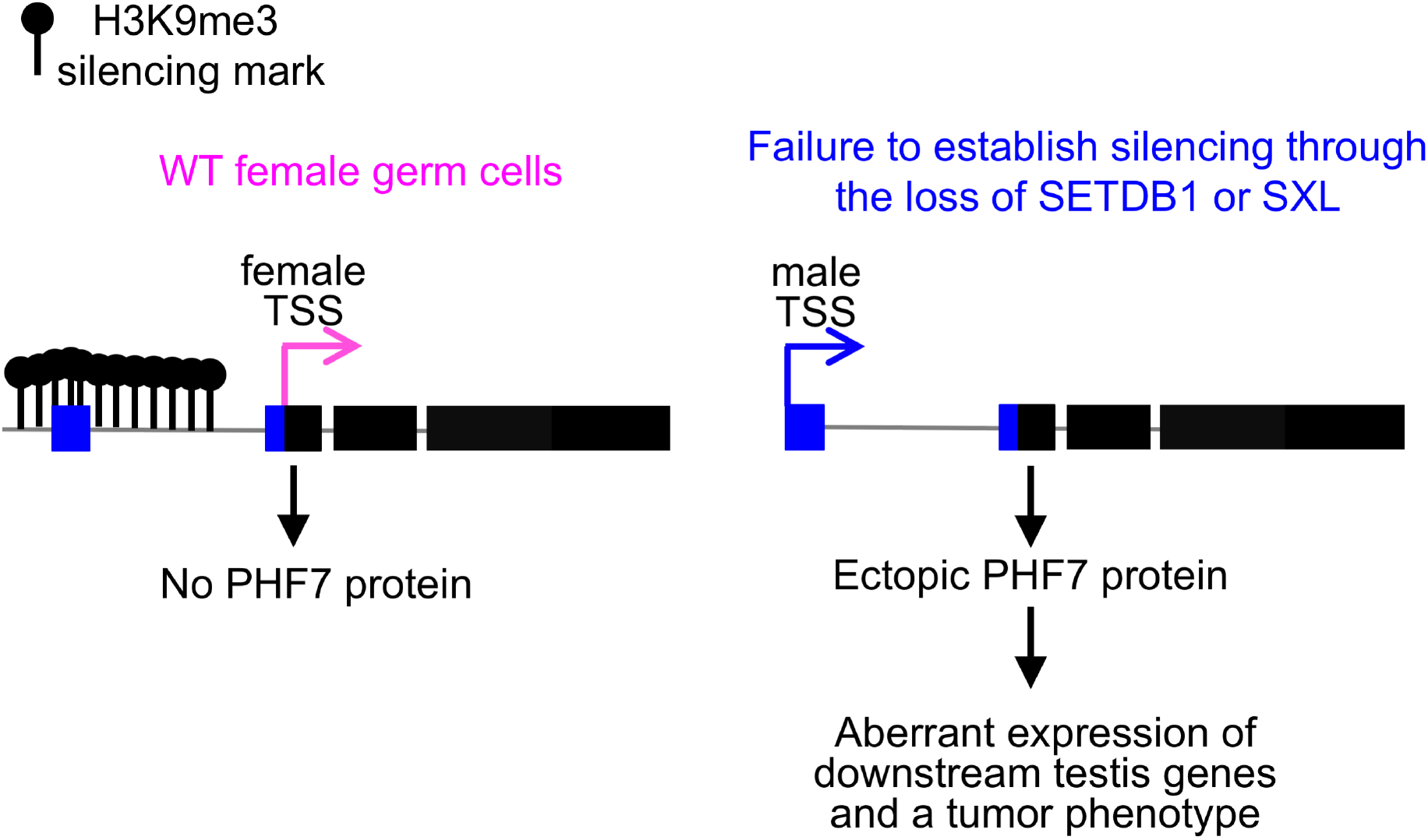
Schematic summary of discrete facultative heterochromatin island assembly at the *phf7* locus. In female germ cells, SETDB1, together with SXL, directs assembly of a highly localized H3K9me3 domain around the testis-specific TSS. In germ cells lacking SETDB1 or SXL protein, the dissolution of the H3K9me3 domain correlates with ectopic testis-specific *phf7-RC* transcription and PHF7 protein expression. Ectopic PHF7 protein activity leads to activation of downstream testis genes and a tumor phenotype.

Prior studies have established a role for SETDB1 in germline Piwi-interacting small RNA (piRNA) biogenesis and TE silencing (Rangan et al., 2011; Sienski et al., 2015; Yu et al., 2015). However, piRNAs are unlikely to contribute to sexual identity maintenance as mutations that specifically interfere with piRNA production, such as *rhino*, do not cause defects in germ cell differentiation (Klattenhoff et al., 2009; Mohn et al., 2014; Volpe et al., 2001; Zhang et al., 2014). Additionally, analysis of differential transcription also shows no masculinization of the gene expression program [(Mohn et al., 2014); data not shown]. Thus, the means by which SETDB1 methylates chromatin at testis genes is likely to be mechanistically different from what has been described for piRNA-guided H3K9me3 deposition on TEs.

We find that H3K9me3 accumulation at many of these genes, including *phf7*, is dependent on the presence of SXL protein. Thus, our studies suggest that SXL is required for female-specific SETDB1 function. SXL encodes an RNA binding protein known to regulate its target genes at the posttranscriptional levels (Salz and Erickson, 2010). SXL control may therefore be indirect. However, studies in mammalian cells suggest that proteins with RNA binding motifs are important for H3K9me3 repression (Becker et al., 2017; Thompson et al., 2015), raising the tantalizing possibility that SXL might play a more direct role in governing testis gene silencing. Further studies will be necessary to clarify how the sex determination pathway feeds into the heterochromatin pathway.

*phf7* stands out among the cohort of genes regulated by facultative heterochromatin because of its pivotal role in controlling germ cell sexual identity (Shapiro-Kulnane et al., 2015; Yang et al., 2012). Because ectopic protein expression is sufficient to disrupt female fate, tight control of *phf7* expression is essential. *phf7* regulation is complex, employing a mechanism that includes alternative promoter usage and TSS selection. We report here that SETDB1/H3K9me3 plays a critical role in controlling *phf7* transcription. In female germ cells, H3K9me3 accumulation is restricted to the region surrounding the silent testis-specific transcription start site. Dissolution of the H3K9me3 marks via loss of SXL or SETDB1 protein is correlated with transcription from the upstream testis-specific site and ectopic protein expression, demonstrating the functional importance of this histone modification. Together, these studies suggest that maintaining the testis *phf7* promoter region in an inaccessible state is integral to securing female germ cell fate.

Although the loss of H3K9me3 pathway members in female germ cells leads to global derepression of testis genes, our integrative analysis identified only 21 SETDB1/H3K9me3 regulated genes. Given that one of these genes is *phf7* and that ectopic PHF7 is sufficient to destabilize female fate (Shapiro-Kulnane et al., 2015; Yang et al., 2012), it is likely that inappropriate activation of a substantial number of testis genes is a direct consequence of ectopic PHF7 protein expression. How PHF7 is able to promote testis gene transcription is not yet clear. PHF7 is a PHD-finger protein that preferentially binds to H3K4me2 (Yang et al., 2012), a mark associated with poised, but inactive genes and linked to epigenetic memory (Pekowska et al., 2010; Pinskaya and Morillon, 2009; Zhang et al., 2012). Thus, one simple model is that ectopic PHF7 binds to H3K4me2 marked testis genes to tag them for transcriptional activation.

It will be interesting to explore whether any of the other 20 SETDB1/H3K9me3 regulated genes also have reprogramming activity. In fact, ectopic fate-changing activity has already been described for the homeobox transcription factor Lim1 in the eye-antenna imaginal disc (Roignant et al., 2010). However, whether Lim1 has a similar function in germ cells is not yet known. Intriguingly, protein prediction programs identify three of the uncharacterized testis-specific genes as E3 ligases (Gene2Function.org) (Hu et al., 2017a). SkpE is a member of the SKP1 gene family, which are components of the Skp1-Cullin-F-box type ubiquitin ligase. CG12477 is a RING finger domain protein, most of which are believed to have ubiquitin E3 ligases activity. CG42299 is closely related to the human small ubiquitin-like modifier (SUMO) E3 ligase NSMCE2. Given studies that have linked E3 ligases to the regulation of chromatin remodeling (Dubiel et al., 2018; Wotton et al., 2017), it is tempting to speculate that ectopic expression of one or more of these testis-specific E3 ligases will be sufficient to alter cell fate. Future studies focused on this diverse group of SETDB1/H3K9me3 regulated genes and their role in reprogramming may reveal the multiple layers of regulation required to secure sexual identity.

The SETDB1-mediated mechanism for maintaining sexual identity we have uncovered may not be restricted to germ cells. Recent studies have established that the preservation of sexual identity is essential in the adult somatic gut and gonadal cells for tissue homeostasis (Grmai et al., 2018; Hudry et al., 2016; Ma et al., 2016; Ma et al., 2014; Regan et al., 2016). Furthermore, the finding that loss of HP1a in adult neurons leads to masculinization of the neural circuitry and male specific behaviors (Ito et al., 2012) suggests a connection between female identity maintenance and H3K9me3 chromatin. Thus, we speculate that SETDB1 is more broadly involved in maintaining female identity.

Our studies highlight an emerging role for H3K9me3 chromatin in cell fate maintenance (Becker et al., 2016). In the fission yeast *S. pombe*, discrete facultative heterochromatin islands assemble at meiotic genes that are maintained in a silent state during vegetative growth (Nakayama et al., 2001; Zofall et al., 2012). Although less well understood, examples in mammalian cells indicate a role for SETDB1 in lineage-specific gene silencing (Du et al., 2018; Jiang et al., 2017; Koide et al., 2016; Schultz et al., 2002; Tan et al., 2012). Thus, silencing by SETDB1 controlled H3K9 methylation may be a widespread strategy for securing cell fate. Interestingly, H3K9me3 chromatin impedes the reprogramming of somatic cells into pluripotent stem cells (iPSCs). Conversion efficiency is improved by depletion of SETDB1 (Chen et al., 2013; Soufi et al., 2012; Sridharan et al., 2013). If erasure of H3K9me3 via depletion of SETDB1 alters the sexually dimorphic gene expression profile in reprogrammed cells, as it does in Drosophila germ cells, the resulting gene expression differences may cause stem cell dysfunction, limiting their therapeutic utility.

## Materials and Methods

### Drosophila stocks and culture conditions

Fly strains were kept on standard medium at 25°C unless otherwise noted. Knockdown studies were carried out with the following lines generated by the *Drosophila* Transgenic RNAi Project (Hu et al., 2017b; Ni et al., 2011; Perkins et al., 2015): *setdb1-P{TRiP.HMS00112}* (BDSC #34803, RRID: BDSC_34803), *Su(var)205/HP1a-P{TRiP.GL00531}* (BDSC #36792, RRID: BDSC_36792), and *wde-P{TRiP.HMS00205}* (BDSC #33339, BDSC_33339). Different conditions were used to maximize the penetrance of the germ cell tumor phenotype. For knockdown of *setdb1* and *HP1a* the *nos-Gal4;bam-Gal80* driver was used (Matias et al., 2015), crosses were set up at 29°C and adults were aged 3-5 days prior to gonad dissection. For *wde* knockdown the *nos-Gal4* driver was used (BDSC #4937, RRID: BDSC_4937) (van Doren et al., 1998), crosses were set up at 18°C and adults were transferred to 29°C for 2 days prior to gonad dissection. The following Drosophila stocks were also used in this study: *snf^148^* (BDSC #7398) (Nagengast et al., 2003) and *HA-setdb1* (Seum et al., 2007). Wild-type ovaries are either sibling controls, or *y^1^ w^1^* (BDSC #1495, RRID: BDSC_1495).

### Immunofluorescence and image analysis

*Drosophila* gonads were fixed and stained according to our previously published procedures (Chau et al., 2009). The following primary antibodies were used: mouse α-Spectrin (1:100, Developmental Studies Hybridoma Bank [DSHB] 3A9 RRID: AB_528473], rabbit α-H3K9me3 (1:1,000, Active Motif Cat# 39162, RRID: AB_2532132), rat α-HA (1:500, Roche Cat# 11867423001, RRID: AB_390919), mouse α-Sxl (1:100, DSHB M18 RRID: AB_528464), rabbit α-Vasa (1:2,000, a gift from the Rangan lab), and rat α-Vasa (1:100, DSHB RRID: AB_760351). Staining was detected by FITC (1:200, Jackson ImmnoResearch Labs) or Alexa Fluor 555 (1:200, Life Technologies) conjugated species appropriate secondary antibodies. TO-PRO-3 Iodide (Fisher, Cat# T3605) was used to stain DNA. Images were taken on a Leica SP8 confocal and compiled with Gnu Image Manipulation Program (GIMP) and Microsoft PowerPoint. All staining experiments were replicated at least two times. The “n” in the figure legends represents the number of germaria scored from a single staining experiment.

### qRT-PCR and data analysis

RNA was extracted from dissected ovaries using TRIzol (Invitrogen, Cat# 15596026) and DNase (Promega, Cat# M6101). Quantity and quality were measured using a NanoDrop spectrophotometer. cDNA was generated by reverse transcription using the SuperScript First-Strand Synthesis System Kit (Invitrogen, Cat# 11904018) using random hexamers. Quantatative real-time PCR was performed using Power SYBR Green PCR Master Mix (ThermoFisher, Cat# 4367659) with the Applied Biosystems 7300 Real Time PCR system. PCR steps were as follows: 95°C for 10 minutes followed by 40 cycles of 95°C for 15 seconds and 60°C for 1 minute. Melt curves were generated with the following parameters: 95°C for 15 seconds, 60°C for 1 minutes, 95°C for 15 seconds, and 60°C for 15 seconds. Measurements were taken in biological triplicate with two technical replicates. The *phf7-RC* levels were normalized to the total amount *phf7*. Relative transcript amount were calculated using the 2-ΔΔCt method (Livak and Schmittgen, 2001). Primer sequences for measuring the total *phf7* and *phf7-RC* levels were: for total *phf7*, forward GAGCTGATCTTCGGCACTGT and reverse GCTTCGATGTCCTCCTTGAG; for *phf7-RC* forward AGTTCGGGAATTCAACGCTT and reverse GAGATAGCCCTGCAGCCA.

### RNA-seq and data analysis

Total RNA was extracted from dissected ovaries using standard TRIzol (Invitrogen, Cat# 15596026) methods. RNA quality was assessed with Qubit and Agilent Bioanalyzer. Libraries were generated using the Illumina TruSeq Stranded Total RNA kit (Cat# 20020599). Sequencing was completed on 2 biological replicates of each genotype with the Illumina HiSeq 2500 v2 with 100bp paired end reads. Sequencing reads were aligned to the *Drosophila* genome (UCSC dm6) using TopHat (2.1.0) (Trapnell et al., 2009). Differential analysis was completed using CuffDiff (2.2.1) (Trapnell et al., 2012). Genes were considering differentially expressed if they exhibited a two-fold or higher change relative to wild-type with a False Discovery Rate (FDR) <5%. Genes that were expressed in mutant (FPKM ≥ 1) but not expressed in wild-type ovaries (FPKM<1) were called ectopic.

Testes genes were identified by interrogating the published mRNA-seq data sets GSE86974 (Shan et al., 2017), as above. Genes whose expression levels were two-fold or higher in *bgcn* mutant testis relative to wild-type testis were called “early-stage testis genes”. Genes whose expression levels were two-fold or higher in wild-type testis relative to *bgcn* mutant testis were called “late-stage testis genes”. Genes that were not differentially expressed but expressed in testis (FPKM ≥ 1) were called simply “testis genes”. The RNA-seq data on Fly Atlas (Leader et al., 2018) was used to identify genes with testis-specific isoforms, i.e. no expression in any other adult tissue.

Heat maps were generated using the CummeRbund R package (2.18.0) (Trapnell et al., 2012). Screen shots are from Integrated Genome Viewer (IGV). To account for the differences in sequencing depth when creating IGV screenshots, the processed RNAseq alignment files were scaled to the number of reads in the wild-type file. This was done with Deeptools bigwigCompare using the scale Factors parameter with a bin size of 5.

### ChIP-seq and data analysis

For each chromatin immunoprecipitation, 400 pairs of *Drosophila* ovaries were dissected in PBS plus protease inhibitors. Tissues were fixed in 1.8% formaldehyde for 10 minutes at room temperature and quenched with 225 mM glycine for 5 minutes. Tissues were washed twice and stored at -80°C for downstream applications. Samples were lysed using the protocol from Lee et al, 2006. Tissue was placed in lysis buffer 1 (140 mM HEPES pH 7.5, 200 mM NaCl, 1 mM EDTA, 10% glycerol, 0.5% NP-40, 0.25% Triton X-100), homogenized using sterile beads, and rocked at 4°C for 10 minutes. Tissue was then washed 10 minutes at 4°C in lysis buffer 2 (10 mM Tris pH 8, 200 mM NaCl, 1 mM EDTA, 0.5 mM EGTA). Tissues were then placed in 1.5 mL lysis buffer 3 (10 mM Tris pH 8, 100 mM NaCl, 1 mM EDTA, 0.5 mM EGTA, 0.1% Na-deoxycholate, 0.5% N-lauroylsarcosine). All buffers were supplemented with protease inhibitors. Chromatin was sheared to 200-700 base pairs using the QSONICA sonicator (Q800R). The chromatin lysate was incubated overnight at 4°C with H3K9me3 antibodies pre-bound to magnetic beads. The beads were prepared as follows: 25 μl Protein A and 25 μl Protein G Dynabeads (Invitrogen, Cat# 10002D and 10004D) per sample were washed twice with ChIP blocking buffer (0.5% Tween 20, 5 mg/mL BSA), then blocked by rocking at 4°C for 1 hour in ChIP blocking buffer, and then conjugated to 5 μg H3K9me3 antibody (Abcam Cat# 8898 RRID: AB_306848) by rocking in new ChIP blocking buffer at 4°C for 1 hour. Following immunoprecipitation, the samples were washed 6 times with ChIP-RIPA buffer (10 mM Tris-HCl pH 8, 1 mM EDTA, 140 mM NaCl, 1% Triton X-100, 0.1% SDS, 0.1% Na-Deoxycholate), 2 times with ChIP-RIPA/500 buffer (ChIP-RIPA + 500 mM NaCl), 2 times with ChIP-LiCl buffer (10 mM Tris-HCl pH 8, 1 mM EDTA, 250 mM LiCl, 0.5% NP-40, 0.5% Na-deoxycholate), and twice with TE buffer. DNA was eluted from beads with 50 μl elution buffer (10 mM Tris-HCl pH 8, 5 mM EDTA, 300 mM NaCl, 0.1% SDS) and reverse crosslinked at 65°C for 6 hours. Beads were spun down and eluted DNA was transferred to a new tube and extracted using phenol-chloroform extraction. All buffers were supplemented with protease inhibitors.

ChIP sequencing libraries were prepared using the Rubicon Genomics Library Prep Kit (Cat# R440406) with 16 amplification cycles. DNA was cleaned and assessed for quality with Qubit and Agilent Bioanalyzer. Sequencing was completed on 2 biological replicates of each genotype with the Illumina HiSeq 2500 v2 with 50 bp single end reads.

H3K9me3 reads were aligned to the *Drosophila* genome (UCSC dm6) using bowtie2 (2.2.6) (Langmead and Salzberg, 2012), and duplicate reads were removed with samtools (1.3) (Li et al., 2009). Peaks were called with MACS (2.1.20150731) using the broadpeaks option with all other paramaters set to default (Zhang et al., 2008). Differential peak analysis on all replicates was completed with the DIFFBIND program (2.4.8), using summits=500 and the DESeq2 package http://bioconductor.org/packages/release/bioc/vignettes/DiffBind/inst/doc/DiffBind.pdf. The mean peak concentration was calculated by normalizing reads to the total library size and subtracting the corresponding input reads. Differential peak fold changes were calculated by subtracting wild-type mean concentrations from mutant mean concentrations. Mutant peaks were considered significantly altered relative to wild-type if they had a False Discovery Rate (FDR) < 5%. The average H3K9me3 deposition on genes in wild-type ovaries was generated with deeptools (2.5.3), using normalized ChIP reads from wild-type ovaries (Ramirez et al., 2016). Screen shots are from Integrated Genome Viewer (IGV).

## Data availability

RNA-seq and ChIP-seq data sets are available from the National Center for Biotechnology Information’s GEO database (http://www.ncbi.nlm.nih.gov/geo/) under accession number GSE109852.

## Acknowledgments

We thank Jean-Rene Huynh, Mark Van Doren, Prash Rangan, the Bloomington *Drosophila* Stock Center and the Iowa Developmental Studies Hybridoma Bank for fly stocks and antibodies; Alex Miron, Neil Molyneaux, Ricky Chan, Ulrich Ness, Olivia Corradin, Dan Factor, Alina Saiakhova for help with the bioinformatic analysis; Heather Broihier, Michelle Longworth, Ron Conlon, Peter Harte and Peter Scacheri for helpful suggestions, discussions and comments on the manuscript; and Jane Heatwole for fly food. This work was supported by the National Institutes of Health, R01GM102141 to HKS and T32GM008056 to AES. Imaging was performed using equipment purchased through NIH S10OD016164.

## Author Contributions

Conceptualization, A.E.S and H.K.S.; Investigation, methodology and analysis A.E.S and L.S.K.; Manuscript writing, reviewing and editing, A.E.S and H.K.S.; Supervision, H.K.S.

---

**Fig. S1.**
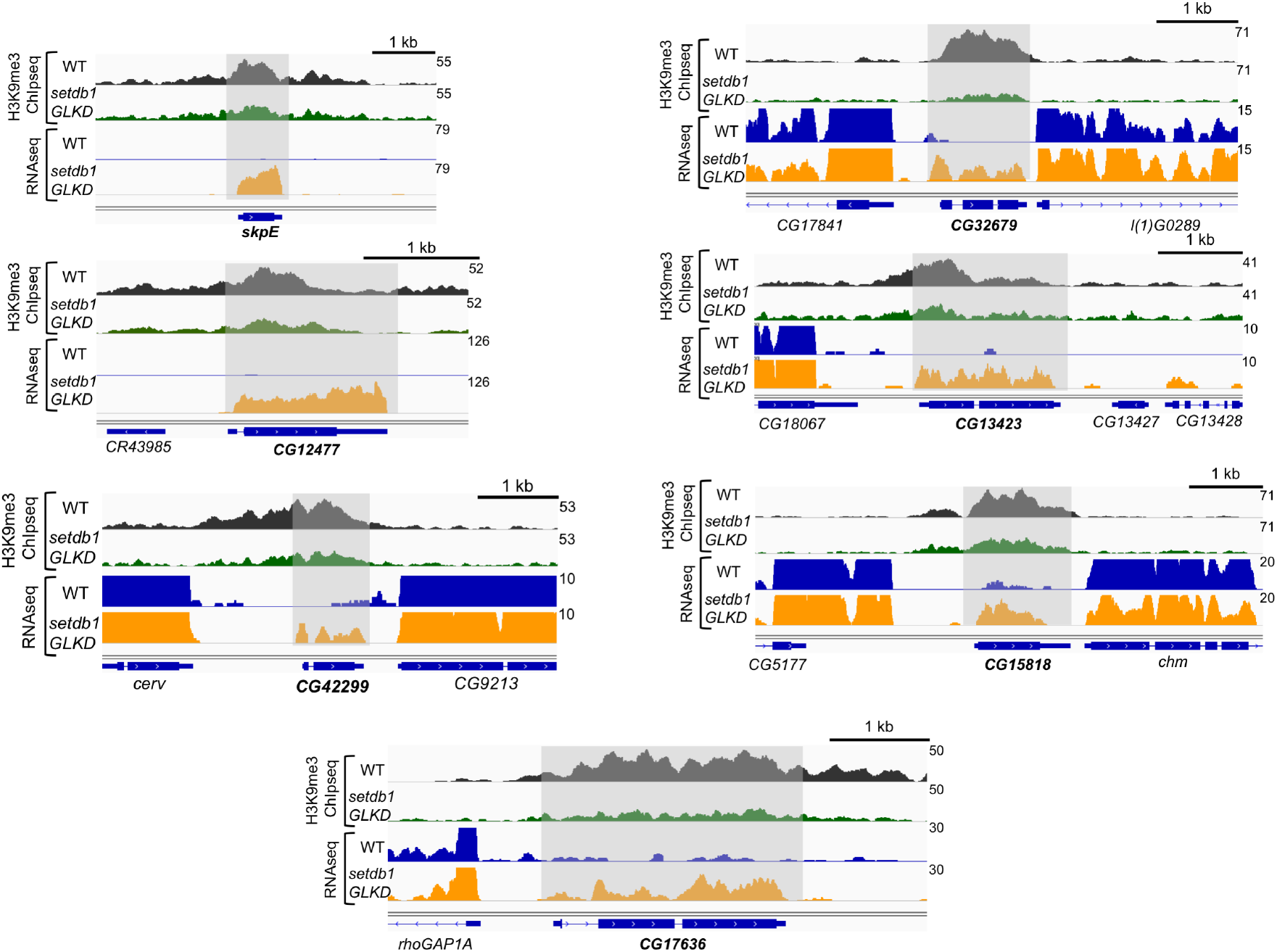
Examples of SETDB1/H3K9me3 regulated genes. Genome browser views of gene neighborhoods illustrate that the H3K9me3 islands present in WT ovaries does not spread to neighboring loci. In each example, shading highlights the gene whose ectopic expression in *setdb1* GLKD is correlated with a decreased H3K9me3 ChIP-seq peak.

**Table S1** Genes upregulated at least 2-fold in *setdb1 GLKD* ovaries.

**Table S2** Genes upregulated at least 2-fold in *wde GLKD* ovaries.

**Table S3** Genes upregulated at least 2-fold in *hp1a GLKD* ovaries.

**Table S4** Genes downregulated at least 2-fold in *setdb1 GLKD* ovaries.

**Table S5** Genes downregulated at least 2-fold in *wde GLKD* ovaries.

**Table S6** Genes downregulated at least 2-fold in *hp1a* GLKD ovaries.

**Table S7** Many of the genes ectopically expressed in *setdb1 GLKD* ovaries are normally expressed in testis. Data from WT testis and *bgcn* mutant testis from (Shan et al., 2017).

**Table S8** Many of the genes ectopically expressed in *wde GLKD* ovaries are normally expressed in testis. Data from WT testis and *bgcn* mutant testis from (Shan et al., 2017).

**Table S9** Many of the genes ectopically expressed in *hp1a GLKD* ovaries are normally expressed in testis. Data from WT testis and *bgcn* mutant testis from (Shan et al., 2017).

**Table S10.** Genomic locations of the 21 SETDB1/H3K9me3 regulated genes.

| gene | Chromosome<br>& sequence location | Cytogenetic<br>map | Distance from<br>nearest<br>neighbor (Mb) |
| --- | --- | --- | --- |
| CG17636 | X:124,370..126,714 [-] | 1A | -- |
| CG34434 | X:5,614,554..5,616,370 [-] | 5A8 | 5.5 |
| Lim1 | X:8,756,539..8,805,804 [-] | 8A5-8B2 | 3.1 |
| CG32679 | X:10,366,402..10,367,472 [-] | 9B12 | 1.6 |
| Rab3-GEF | X:15,089,682..15,113,762 [+] | 13A10-12 | 4.7 |
| CG42299 | X:15,574,947..15,575,683 [-] | 13D | 0.5 |
| CR43299 | X:15,662,289..15,665,707 [-] | 13E | 0.1 |
| SkpE | X:19,823,368..19,824,080 [-] | 18F | 4.2 |
| Phf7 | X:20,160,355..20,165,830 [-] | 19B3-19C1 | 0.3 |
| CG32506 | X:20,449,331..20,467,065 [+] | 19D1 | 0.3 |
| CG12061 | X:23,036,066..23,047,678 [+] | 20F4 | 2.6 |
| CG15818 | 2L:7,410,909..7,412,229 [+] | 27F3 | -- |
| CG15172 | 2L:19,022,417..19,023,646 [+] | 37B9 | 11.6 |
| CG13423 | 2R:20,536,578..20,538,264 [+] | 57A5 | -- |
| CG10440 | 2R:21,560,246..21,576,035 [-] | 57F1-2 | 1.0 |
| MsR1 | 3L:2,322,693..2,345,799 [+] | 62D6 | -- |
| CG12607 | 3L:4,446,573..4,448,072 [+] | 64B6 | 2.1 |
| CG10483 | 3L:5,917,905..5,921,316 [-] | 64F5 | 1.5 |
| CG12477 | 3L:18,721,893..18,723,272 [-] | 75D7 | 12.8 |
| CG42613 | 3R:18,929,196..18,978,088 [+] | 91D5-91E1 | -- |
| CG31202 | 3R:29,928,701..29,930,386 [-] | 99C7 | 10.9 |

